# Sex-specific transgenerational plasticity II: Grandpaternal effects are lineage- and sex-specific in threespined sticklebacks

**DOI:** 10.1101/796995

**Authors:** Jennifer K Hellmann, Erika R Carlson, Alison M Bell

**Author notes:** Corresponding author: Jennifer Hellmann, 505 S Goodwin Ave, Urbana IL 61801.

## Abstract

1. Transgenerational plasticity (TGP) occurs when the environment encountered by one generation (F0) alters the phenotypes of one or more future generations (e.g. F1 and F2). Sex selective TGP, via specific lineages or to only male or female descendants, has been underexplored in natural systems, and may be adaptive if it allows past generations to fine-tune the phenotypes of future generations in response to sex-specific life history strategies.
2. We sought to understand if exposing males to predation risk can influence grandoffspring via sperm in threespined stickleback *(Gasterosteus aculeatus).* We specifically tested the hypothesis that grandparental effects are transmitted in a sex-specific way down the male lineage, from paternal grandfathers to F2 males.
3. We reared F1 offspring of unexposed and predator-exposed F0 males under ‘control’ conditions and used them to generate F2s with control grandfathers, a predator-exposed maternal grandfather (i.e., predator-exposed F0 males to F1 daughters to F2 offspring), a predator-exposed paternal grandfather (i.e., predator-exposed F0 males to F1 sons to F2 offspring), or two predator-exposed grandfathers. We then assayed male and female F2s for a variety of traits related to antipredator defense.
4. We found little evidence that transgenerational effects were mediated to only male descendants via the paternal lineage. Instead, grandpaternal effects depended on lineage and were mediated largely across sexes, from F1 males to F2 females and from F1 females to F2 males. When their paternal grandfather was exposed to predation risk, female F2s were heavier and showed a reduced change in behavior in response to a simulated predator attack relative to offspring of control, unexposed grandparents. In contrast, male F2s showed reduced antipredator behavior when their maternal grandfather was exposed to predation risk. However, these patterns were only evident when one grandfather, but not both grandfathers, was exposed to predation risk, suggesting the potential for non-additive interactions across lineages.
5. If sex-specific and lineage effects are common, then grandparental effects are likely underestimated in the literature. These results draw attention to the importance of sex-selective inheritance of environmental effects and raise new questions about the proximate and ultimate causes of selective transmission across generations.

## Introduction

Transgenerational plasticity (TGP, environmental parental effects) occurs when the environment experienced by a parent influences the phenotype of one or more future generations. Although the importance of maternal environments for offspring is well-appreciated (Bonduriansky & Day 2008; Sheriff *et al.* 2017), recent studies are showing the importance of paternal experiences such as stress (Rodgers *et al.* 2013; Dias & Ressler 2014; Gapp *et al.* 2014), diet (Ng *et al.* 2010; de Castro Barbosa *et al.* 2016), and immune priming (Beemelmanns & Roth 2016) for offspring, and that paternal effects can be caused by epigenetic changes to sperm (e.g., small RNAs (Rodgers *et al.* 2015; Immler 2018)). In some cases, parental environments influence the phenotypes of the F1 generation but their effects do not persist into future (e.g. F2) generations (Remy 2010; Wibowo *et al.* 2016; Zhou *et al.* 2018). However, there is also evidence that the effects of environments experienced by one generation (F0) can persist for multiple generations (Kaati, Bygren & Edvinsson 2002; Dias & Ressler 2014; Gapp *et al.* 2014; Cropley *et al.* 2016; Beemelmanns & Roth 2017), even when offspring (F1s) are raised under ‘control’ conditions, i.e. in the absence of the cue that triggered a response in the F0 generation (grandparental effects).

Although multigenerational inheritance can be non-selective (i.e. all grandoffspring are equally affected by their grandparents’ environments; (Bell & Hellmann 2019)), epigenetic changes may also persist selectively across generations in only a subset of individuals. Indeed there is some evidence that transgenerational effects can persist in a lineage-specific (to F2s via either the paternal or maternal lineage) and/or sex-specific (to only male or female F2s) fashion through multiple generations (Dunn & Bale 2011; Lock 2012; Saavedra-Rodríguez & Feig 2013; Bygren *et al.* 2014; Moisiadis *et al.* 2017; Wylde *et al.* 2019). In humans, for instance, grandsons are influenced by the diet of their paternal grandfather while grand-daughters are influenced by the diet of their paternal grandmother (Pembrey *et al.* 2006). Multigenerational studies simultaneously examining both lineage and sex-specific effects have been conducted almost exclusively in mammals (but see (Emborski & Mikheyev 2019)), where mechanisms such as sex-specific placental function and provisioning can generate sex-specific effects in the F1 generation (Bale 2011; Glover & Hill 2012; Yun, Lee & Kim 2016) that might affect the ways in which these effects are transmitted to the F2 generation. Sex-specific and lineage-specific effects in externally-fertilizing organisms or organisms that lack parental care are not well characterized.

Our understanding of lineage and sex-specific effects is limited because they are difficult to study, as they require measuring traits in both male and female F2s and tracking effects through both the maternal or paternal lineage (rather than comparing F2s with control grandparents to F2s with two or four experimental grandparents). By measuring traits in both sexes and tracking lineage, we can learn whether effects are passed only via either the male or female line (e.g., F0 males to F1 males to F2 males) and whether there are interactive effects across lineages. For example, receiving cues from both the maternal and paternal grandfather may result in different traits or more extreme trait values than receiving cues from only the maternal (F0 males to F1 females to F2s) or paternal (F0 males to F1 males to F2s) grandfather.

Sex-specific and lineage-specific effects may have adaptive significance if they can allow past generations to fine-tune the phenotypes of future generations in response to sex-specific life history strategies or sex differences in the costs and benefits of attending to grandparental cues, which might contain outdated and inaccurate information about the environment. However, recent theoretical work has shown that nonadaptive sex-specific parental effects may arise due to sexual conflict, especially in systems with strong sexual selection (Burke, Nakagawa & Bonduriansky 2019).

In a recent study (Hellmann *et al.* in review) we demonstrated that paternal experience with predation risk prior to fertilization produced different phenotypes in F1 sons compared to F1 daughters in threespined stickleback *(Gasterosteus aculeatus).* Here, we track these effects into the next generation to also understand the extent to which sex-specific or lineage-specific effects are present in the F2 generation, which could arise, for example, because transgenerational effects passed through male F1s of predator-exposed fathers differ from transgenerational effects passed through female F1s of predator-exposed fathers. We predicted that cues from F0 males would be transmitted to F1 males to F2 males, but not to F2 females (statistically significant paternal lineage by F2 sex interaction). This could arise because there are a variety of male-specific reproductive traits, such as bright nuptial coloration and conspicuous defense, courtship, and parenting behavior (Bell & Foster 1994), that increase the vulnerability of male sticklebacks to predation risk (Candolin 1998; Johnson & Candolin 2017) and therefore, increase the potential adaptive benefits for male, but not female, sticklebacks to attend to grandpaternal cues of predation risk. Further, regardless of whether epigenetic changes are adaptive or non-adaptive, transmission down the paternal lineage to male descendants could arise via epigenetic changes that are linked to the Y chromosome, such as stable inheritance of methylation (Zhang *et al.* 2016), sex chromosome imprinting (Lemos *et al.* 2013), or Y chromosome polymorphisms (Francisco & Lemos 2014).

To understand the extent to which cues of predation risk in the F0 generation altered the phenotypes of the F2 generation, we reared sons and daughters of control and predator-exposed fathers under ‘control’ conditions (see Part I, (Hellmann *et al.* in review)) and used them to generate F2s with control grandfathers (i.e., offspring of F1 daughters and sons of control males), a predator-exposed maternal grandfather (i.e., offspring of F1 daughters of predator-exposed males and F1 sons of control males), a predator-exposed paternal grandfather (i.e., offspring of F1 daughters of control males and F1 sons of predator-exposed males), or two predator-exposed grandfathers (i.e., offspring of F1 daughters and sons of predator-exposed males; Figure 1). For paternal effects, only phenotypic changes in the F2 generation and beyond are considered truly transgenerational, as any environment experienced by F0 males simultaneously exposes both the father (F0) and his germline (future F1) (Heard & Martienssen 2014). We then assayed male and female F2s to determine if grandpaternal predation exposure induced traits related to antipredator defense (Bell, McGhee & Stein 2016; Sheriff *et al.* 2017), including increased antipredator behaviors in an open field assay as well as altered stress-induced cortisol levels and body size.

**Figure 1:**
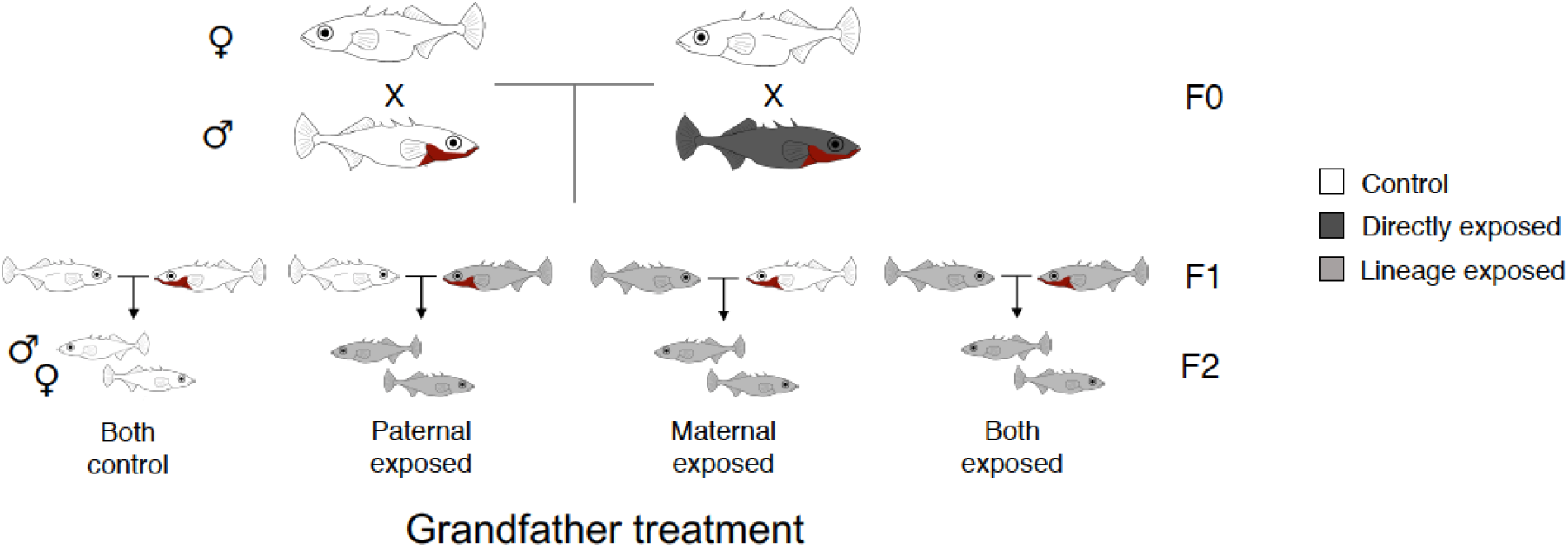
Wild-caught males in the F0 generation were either left unexposed as controls (white) or directly exposed to predation risk (dark grey) and their sperm was used to fertilize the eggs of unexposed, wild-caught females using in vitro fertilization. The F1 generation was reared in the absence of predation risk and used to generate the F2 generation. For example, F1 sons of predator-exposed fathers were mated to F1 daughters of control fathers to generate F2s with a predator-exposed paternal grandfather and F1 daughters of predator-exposed fathers were mated to F1 sons of control fathers to generate F2s with a predator-exposed maternal grandfather. Similarly, F1 daughters of predator-exposed fathers were mated to F1 sons of predator-exposed fathers to generate F2s with both grandfathers exposed to predation risk. Light grey indicates F1s/F2s whose lineage was exposed to predation risk (i.e. their parents or grandparents experienced predation risk). Juvenile F2s were then assayed for a variety of traits.

## Methods

### Housing conditions

In June-September 2016, adult, sexually mature threespined sticklebacks were collected from Putah Creek, a freshwater stream in northern California and shipped to the University of Illinois at Urbana-Champaign. This population has piscivorous predators, including the prickly sculpin (*Cottus asper*). The F0 generation was maintained on a summer photoperiod schedule (16 L: 8D) at 21° ± 1°C and fed ad libitum daily with a mix of frozen bloodworms (*Chironomus* spp.), brine shrimp (Artemia spp.) Mysis shrimp, and Cyclop-eeze. The F0, F1, and F2 generation were all housed on recirculating, temperature-controlled flowthrough racks, with particular, biological, and UV filters; this allowed for conditions to be standardized among tanks during nesting, egg incubation and fry rearing.

To simulate breeding conditions, where males defend nesting territories while females shoal together, F0 females were housed in groups of n=10 fish per tank while F0 males were isolated in a 26.5L tanks (36L × 33W × 24H cm) with nesting materials; once F0 males had successfully built a nest, they were exposed to a clay model sculpin (21cm long) 6 times over 11 days or left undisturbed during an equivalent time frame (n=16 F0 males total). This predator exposure regimen was designed to mimic conditions that males experience when they move into shallow habitats to nest. We elected to expose males to a relatively short stressor in order to minimize the potential for males to habituate to the model predator (Dellinger *et al.* 2018) and to minimize the possibility of influencing sperm quality; although stickleback males produce sperm in the beginning of the breeding season (Borg 1982), it is possible that a prolonged stressor such as simulated predation risk could influence sperm production and/or quality.

The day after the last exposure, F1 offspring were generated via *in vitro* fertilization using a split-clutch design: an unexposed, wild-caught female’s clutch was split and fertilized by both a control and predator-exposed male. We incubated fertilized F1 eggs in a cup with a mesh bottom placed above an air bubbler and fry were reared until adulthood in 37.9 L (53L × 33W × 24H cm) tanks, with each half-clutch housed in a separate tank (densities ranging from 4-28 individuals per tank). The F1 generation was fed newly hatched brine shrimp for 2 months before transitioning to the same mix of frozen bloodworms, brine shrimp, Mysis shrimp, and Cyclop-eeze noted above. The F1 generation was switched from a summer photoperiod schedule (16 L: 8D) at 21° ± 1°C to a winter light schedule (8 L: 16 D) at 20° ± 1°C at the end of the breeding season (December 2016). The F1s used in this experiment were not used in any behavioral assays nor exposed to predation risk (see Hellmann *et al.* (in review) for more details-F1s used in this experiment were siblings of the F1s with control mothers used in Hellmann *et al.* (in review)).

Once the F1 generation neared reproductive maturity (May 2017), the F1 generation was switched from a winter photoperiod (8 L: 16 D at 20° ± 1°C) to a summer photoperiod schedule (16 L: 8D) at 21° ± 1°C. From August – October 2017, we bred male and females F1s to generate the F2 generation. We isolated adult F1 males in 26.5L tanks (36L × 33W × 24H cm); each tank contained two plastic plants, a sandbox, a clay pot, and algae for nest building. Males were left undisturbed until they had completed their nest, at which point we euthanized the male to obtain sperm. F1 females were maintained in their home tanks (to mimic shoaling in the wild) and fed twice per day to encourage egg production; once females were gravid, we gently squeezed their abdomen to obtain eggs. We used a split-clutch design to generate four grandparental treatment groups. Each F1 females’ eggs were fertilized by F1 sons of control and predator-exposed fathers; similarly, each F1 male sired eggs from F1 daughters of control and predator-exposed fathers (Figure 1). To avoid inbreeding, mated F1s had different F0 mothers and fathers. We successfully generated 32 clutches of half-siblings: F2s with control grandfathers (n=8 clutches), predator-exposed paternal grandfather (n=8 clutches), predator-exposed maternal grandfather (n=8 clutches), and two predator-exposed grandfathers (n=8 clutches). Clutch failure and size did not vary with experimental treatment in either the F1 or F2 generation (supplementary material). In total, the F2 generation used in the assays below were descendants of n = 16 F0 grandfathers and n = 19 F1 fathers (range of n = 7-11 F2s tested per clutch, plus one clutch where only n =2 F2s were tested).

As with the F1 generation, we incubated fertilized eggs with an air bubbler and fry were reared in 37.9 L (53L × 33W × 24H cm) tanks, with each half-clutch housed in a separate tank (mean 18.7 individuals per tank, range: 6-35). Offspring were switched to a winter light schedule (8 L: 16 D) at least one month prior to when assays were conducted. As with the F1 generation, F2 fry were fed newly hatched brine shrimp for two months before transitioning to the mix of frozen food described above.

Because mothers and fathers did not interact prior to fertilization nor with their offspring postfertilization, our experimental design allowed us to completely isolate TGP mediated via gametes while controlling for mate choice and differential allocation due to partner quality or parental care (Mashoodh *et al.* 2012; Stein & Bell 2014; McGhee *et al.* 2015; Mashoodh *et al.* 2018). Further, by using artificial fertilization, we controlled for the selective failure of males to court or parent successfully under stressful conditions, which may result in differences between control and predator-exposed lineages because of selective breeding of a nonrandom sample of individuals.

### Open field assays

When the F2 generation was 5 months old (mean days post-hatching: 157.9, range 146-169 days), we measured activity/exploration, and antipredator behavior (freezing and evasive swimming) using similar methods described in Hellmann *et al.* (in review). Briefly, the testing arena was a circular pool (150cm diameter) divided into eight peripheral sections with a circular section in the middle (using lines drawn on the base of the pool). Fish were placed in an opaque refuge in the center of the arena with its entrance plugged. After a three minute acclimation period, we removed the plug from the refuge, allowed fish to emerge, and then measured the number of different (exploration) and total (activity) sections visited for three minutes after emergence. Fish that did not emerge 5 minutes after the plug was removed were gently released from the refuge; while offspring who emerged naturally were more active/exploratory than fish who were released (generalized linear model with binomial distribution (emerged or released), with activity/exploration as a fixed effect: Z_249_=−2.39, p=0.01), controlling for emergence time did not alter the significance of the activity/exploration results reported below.

After the 3min period, we simulated a predator attack by quickly moving a clay sculpin toward the experimental fish for approximately 5 seconds and then removing the sculpin from the arena. This attack often elicited evasive (jerky) swimming behavior followed by freezing behavior from the fish (we interpret increased time frozen and evasive swimming as greater antipredator behavior); we measured whether or not fish performed the evasive swimming behavior as well as the latency for the fish to resume movement. After the fish resumed movement, we then again measured the number of different and total sections visited for 3 minutes. If the fish remained frozen for greater than 10 minutes (n=25 fish), we ended the trial and considered activity and exploration after the simulated predation attack to be zero. We assayed n=63 F2s with control grandfathers (n=32 females, n=31 males), n=64 F2s with predator-exposed paternal grandfathers (n=35 females, n=29 males), n=61 F2s with predator-exposed maternal grandfathers (n=29 females, n=32 males), and n=63 F2s with two predator-exposed grandfathers (n=30 females, n=33 males). Assays were scored live, by an observer who was standing at least 1m from the pool.

To measure cortisol in response to the predator attack (Mommer & Bell 2013), we netted the fish from the arena 15 minutes after the simulated predator attack and quickly weighed and measured it (standard length: from the tip of the nose to the base of the caudal fin). We euthanized the fish in MS-222 and drew blood from the tail of the fish using a heparinized microhematocrit tube. We centrifuged blood to separate the plasma (StatSpin CritSpin Microhemocrit centrifuge) and immediately froze the plasma at −80 °C. Because many fish had non-reproductively mature gonads, we dissected fish and identified sex based on the presence of immature testes and ovaries; we confirmed the accuracy of this method and sexed any questionable fish using a genetic marker (Peichel *et al.* 2004).

### Plasma cortisol

To measure circulating cortisol, we followed the manufacture’s protocol (Enzo Life Sciences, Plymouth Meeting, PA, USA). All the plasma samples were prepared in 1:10 steroid displacement reagent solution, then ran with a 1:120 dilution and in duplicate. Slopes of the standard curves and a serial dilution curve (1:20 to 1:320) were parallel (t_6_=1.21, p=0.27), indicating that there was negligible matrix interference contributing to systematic measurement error. The intra-assay coefficients of variation were all within acceptable range (3.8%, 2.9%, 4.4%, 4.7%, 4.8%, 3.8%). We ran common samples of pooled plasma on each plate (in quadruplicate as the first two and last two wells of each plate) to calculate the interassay coefficient of variation (13.9%). Samples with a coefficient of variation greater than 15% (n = 2) were removed from the data set. Due to insufficient amount of blood drawn from some offspring, we sampled n=48 F2s with control grandfathers, n=57 F2s with predator-exposed paternal grandfathers, n=44 F2s with predator-exposed maternal grandfathers, and n=49 F2s with two predator-exposed grandfathers.

### Statistical analysis

For the activity and exploration, we used a principal components analysis (R package factoextra (Kassambara & Mundt 2017)) to combine the activity and exploration (Spearman rank correlation of activity and exploration: ρ=0.91, p<0.001). Behaviors were scaled and centered, and we included activity and exploration before and after the simulated predator attack (two data points per individual). We extracted one principal component with an eigenvalue of 1.75 that captured 87.5% of the variation, with larger values indicating individuals who were more active/explorative.

We used linear mixed effects models to test predictors of activity/exploration, standard length, mass, and stress-induced cortisol (square-root transformed for normalization of residuals), and generalized linear mixed models to test predictors of variation in freezing behavior (negative binomial distribution) and evasive swimming (binomial distribution) (R package lme4 (Bates *et al.* 2015)). We ran separate models for male and female F2s because of our a priori hypotheses regarding male-lineage effects (i.e. that F2 male phenotypes would be altered by the experiences of their paternal grandfather) and to avoid higher order interaction terms (maternal grandfather by paternal grandfather by F2 sex interactions) that are difficult to interpret. All models included fixed effects of maternal and paternal grandfather treatment. The models testing predictors of activity/exploration, emergence/freezing behavior, mass, and cortisol also included standard length. The model testing predictors of standard length included tank density (assessed at 4.5 months) and age (days since hatching). To control for genetic differences due to either maternal or paternal grandfather identity, all models included random effects of mother and father identity nested within maternal and paternal grandfather identity, as well as observer identity for the behavioral data; because there were two observations per individual, the model of activity-exploration included random effects of fish ID nested within maternal and paternal identity nested within maternal and paternal grandfather (respectively). We tested for possible interactions between maternal and paternal grandfather treatment (as well as interactions with observation period for the activity/exploration model); we retained statistically significant interactions. When these interactions were present, we investigated those interactions by rerunning the models with grandfather treatment as a 4-factor variable to determine differences between F2s of control grandparents and F2s with either maternal, paternal, or two experimental grandfathers. Effect sizes for all models were calculated using the anova_stats function in R package sjstats (Lüdecke 2020). We removed three datapoints (one female, two male) from the mass/length dataset to normalize the residuals. Degrees of freedom were estimated using Satterthwaite’s method for general linear mixed models, and residual degrees of freedom are reported for generalized linear mixed models.

### Animal welfare note

All methods, including euthanasia techniques, were approved by Institutional Animal Care and Use Committee of University of Illinois Urbana-Champaign (protocol ID 15077).

## Results

### Female F2s were less responsive to a simulated predator attack when their paternal grandfather, but not both grandfathers, was exposed to predation risk

We sought to test the hypothesis that grandparental effects are transmitted in a sex-specific way down the male lineage, from paternal grandfathers to F2 males. For activity/exploration, we found no evidence of main or interactive effects of maternal or paternal grandfather treatment for male F2s (Table 1; Figure 2). For female F2s, we found that activity/exploration was influenced by a three-way interaction between paternal grandfather treatment, maternal grandfather treatment, and observation period (Table 1). Specifically, female F2s with a predator-exposed paternal grandfather showed a reduced change in activity/exploratory behavior in response to the simulated predator attack compared to female F2s with control grandfathers (paternal GF by observation period interaction: t_122.00_=3.03, p=0.003; Figure 2); however, this pattern was not present for female F2s with either a predator-exposed maternal grandfather (t_122.00_=1.40, p=0.17) or two predator-exposed grandfathers (t_122.00_=0.33, p=0.75; Figure 2). To determine if these effects varied across families with different F0 males, we tested whether model fit was worsened when random effects of paternal/paternal grandfather ID or maternal/maternal grandfather ID were removed. Model fit did not significantly worsen when random effects of paternal/paternal 22 grandfather ID (χ^2^=0.94, p=0.82) or maternal/maternal grandfather ID were removed (χ^2^=5.87, p=0.12), suggesting that these transgenerational effects are consistent across different families.

**Figure 2:**
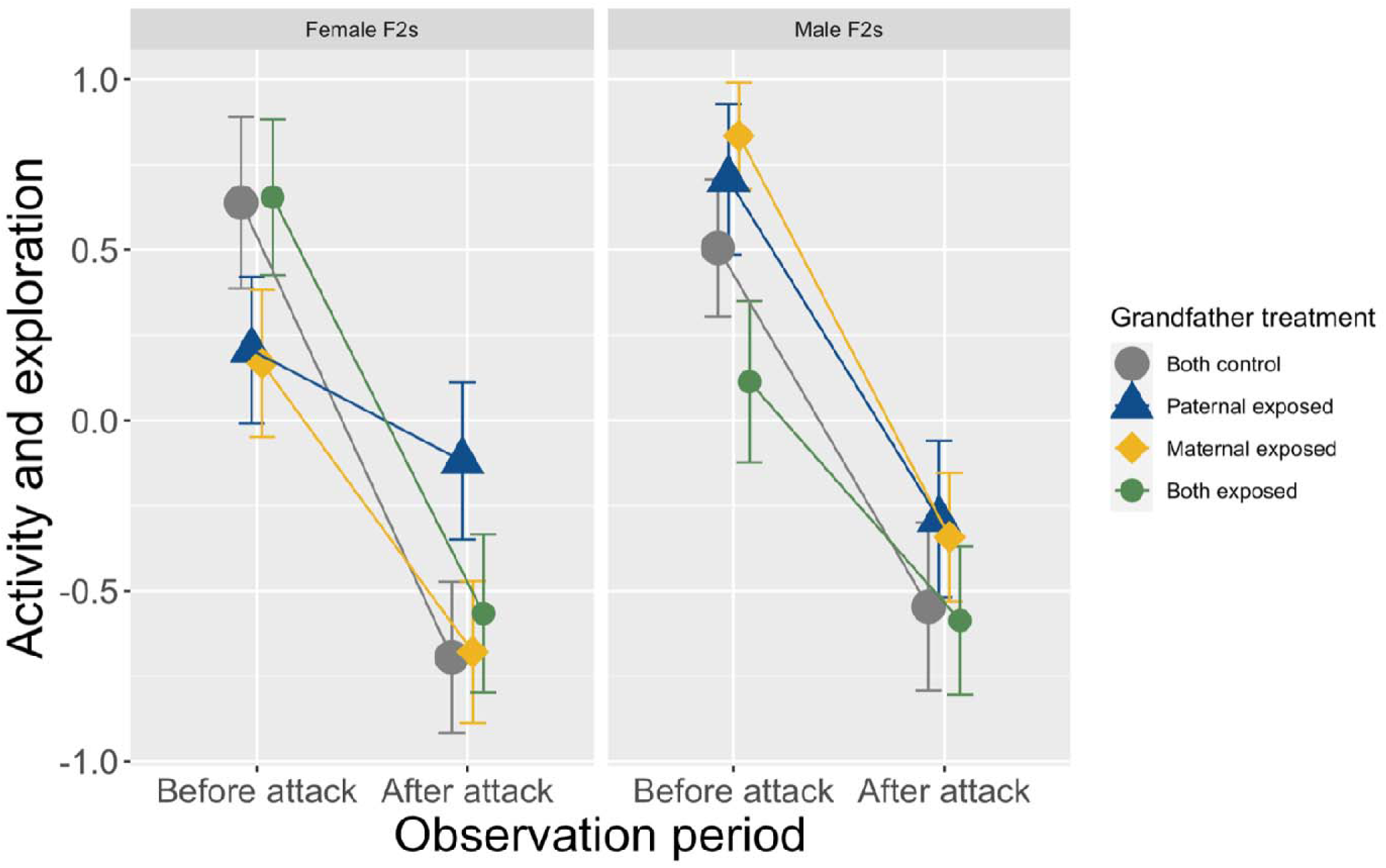
Activity and exploratory behavior (PCA: higher values are more active and exploratory individuals) of female (left) and male (right) F2s before and after the simulated predator attack in the open field assay (means with standard error bars).

**Table 1:** Results of mixed models testing predictors of F2 activity/exploratory behavior (higher values showed more active and exploratory individuals; n=125 males, n=126 females), freezing behavior (n=125 males, n=126 females), and evasive swimming behavior (n=124 males, n=125 females).

|  | Female F2s |  |  |  | Male F2s |  |  |  |
| --- | --- | --- | --- | --- | --- | --- | --- | --- |
|  | Activity and exploratory behavior |  |  |  |  |  |  |  |
|  | *t* | *df* | p | *Cohen's F* | *t* | *df* | p | *Cohen's F* |
| Maternal grandfather treatment | -1.18 | 23.10 | 0.25 | 0.01 | -0.03 | 10.36 | 0.97 | 0.00 |
| Paternal grandfather treatment | -1.88 | 27.00 | 0.07 | 0.02 | -0.48 | 8.58 | 0.64 | 0.03 |
| Observation period | -5.54 | 122.00 | **<0.001** | 0.57 | -7.20 | 123.85 | **<0.001** | 0.48 |
| Standard length | 0.42 | 115.26 | 0.68 | 0.03 | 1.56 | 115.20 | 0.12 | 0.10 |
| Maternal GF * paternal GF | 2.57 | 198.92 | **0.01** | 0.09 | --- | --- | --- | --- |
| Paternal GF * observation period | 3.03 | 122.00 | **0.003** | 0.10 | --- | --- | --- | --- |
| Maternal GF * observation period | 1.40 | 122.00 | 0.17 | 0.06 | --- | --- | --- | --- |
| Maternal GF * paternal GF * obs period | -2.84 | 122.00 | **0.005** | 0.21 | --- | --- | --- | --- |
|  | Freezing behavior |  |  |  |  |  |  |  |
|  | *Z* | *df* | p | *Cohen's F* | *Z* | *df* | p | *Cohen's F* |
| Maternal grandfather treatment | -0.96 | 1, 116 | 0.34 | 0.17 | -1.69 | 1, 114 | 0.09 | 0.08 |
| Paternal grandfather treatment | 0.06 | 1, 116 | 0.95 | 0.05 | -1.68 | 1, 114 | 0.09 | 0.06 |
| Standard length | -0.82 | 1, 116 | 0.41 | 0.12 | -1.50 | 1, 114 | 0.13 | 0.13 |
| Maternal * paternal GF treatment | --- | --- | --- | --- | 1.97 | 1, 114 | **0.049** | 0.16 |
|  | Evasive swimming behavior |  |  |  |  |  |  |  |
|  | *Z* | *df* | p | *Cohen's F* | *Z* | *df* | p | *Cohen's F* |
| Maternal grandfather treatment | -1.38 | 1, 116 | 0.17 | 0.10 | -1.91 | 1, 114 | 0.06 | 0.02 |
| Paternal grandfather treatment | -0.65 | 1, 116 | 0.51 | 0.03 | -0.25 | 1, 114 | 0.80 | 0.16 |
| Standard length | 1.24 | 1, 116 | 0.22 | 0.12 | -0.18 | 1, 114 | 0.86 | 0.02 |
| Maternal * paternal GF treatment | --- | --- | --- | --- | 2.38 | 1, 114 | **0.02** | 0.21 |

### Male F2s showed reduced antipredator behaviors when one grandfather, but not both grandfathers, was exposed to predation risk

For male F2s, there was an interaction between maternal grandfather and paternal grandfather treatment on both freezing behavior and evasive swimming behaviors (Table 1; Figure 3). Specifically, male F2s with a maternal grandfather exposed to predation risk were less likely to perform evasive swimming behaviors (Z_114_=−1.91, p=0.06) after the simulated predator attack relative to offspring of control grandfathers.

**Figure 3:**
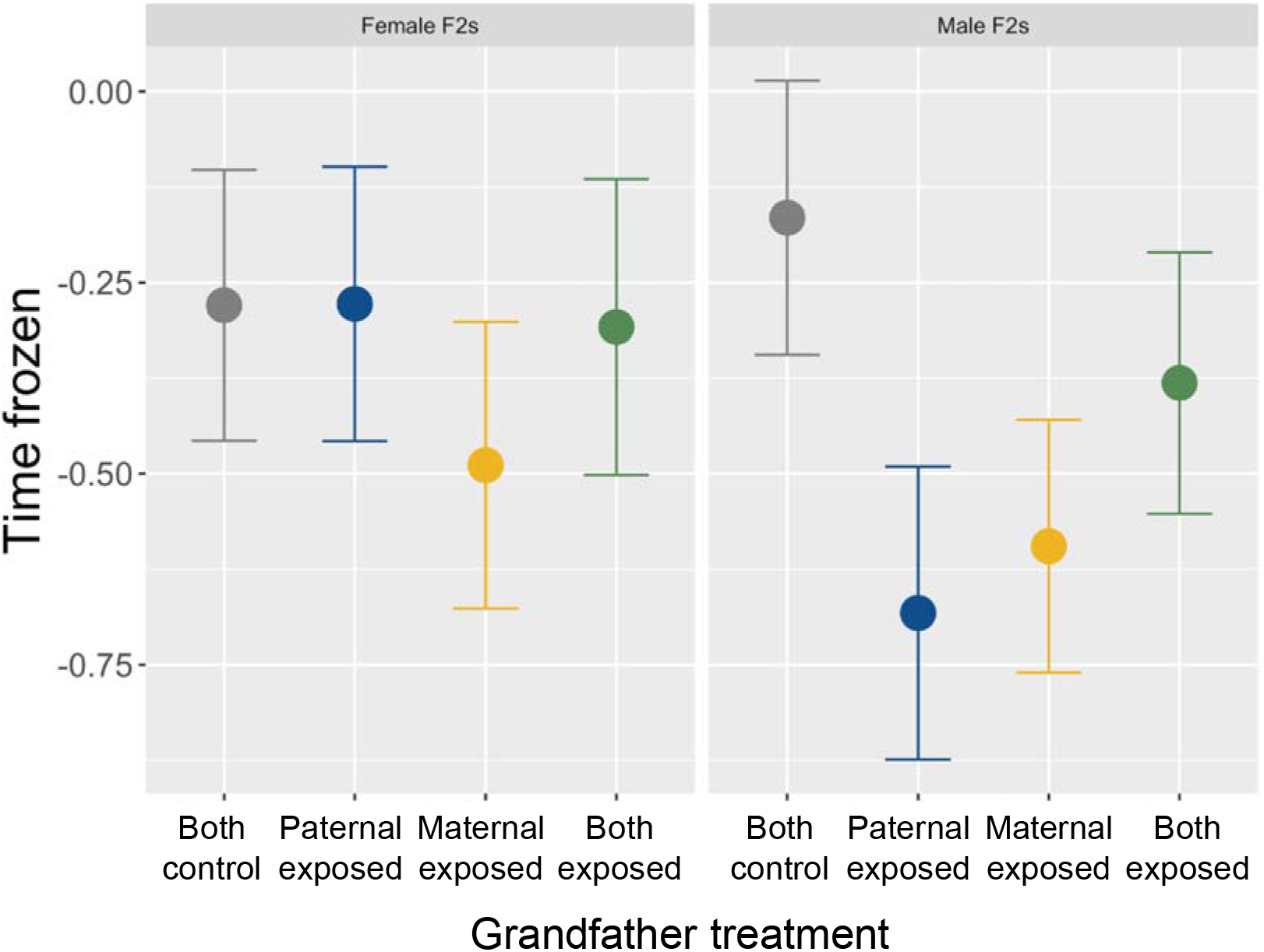
Freezing behavior of female (left; n=126) and male (right; n=125) F2s with two control grandfathers (grey), predator-exposed paternal grandfather (blue), predator-exposed maternal grandfather (yellow), or two predator-exposed grandfathers (green). Shown are the residuals of the regression model with grandpaternal treatment removed, plotted against grandpaternal treatment.

However, there was no detectable difference between male F2s with a control grandfather and male F2s with either a predator-exposed paternal grandfather (Z_114_=−0.25, p=0.80) or two predator-exposed grandfathers (Z_114_=0.91, p=0.36). Similarly, male F2s with a paternal (Z_114_=−1.68, p=0.09) or maternal (Z_114_=−1.69, p=0.09) grandfather exposed to predation risk tended to spend less time frozen after the simulated predator attack relative to offspring of control grandfathers, but this was not true for male F2s with two predator-exposed grandfathers (Z_114_=−0.88, p=0.38; Figure 3). Model fits did not significantly worsen when random effects of paternal/paternal grandfather (jerky swims: χ^2^=0, p=1.00; freezing: χ^2^=0, p=1.00) or maternal/maternal grandfather were removed (jerky swims: χ^2^=1.28, p=0.53; freezing: χ^2^=4.76, p=0.09), suggesting that genetic variation in the F0 and F1 generation does not strongly affect either behavior for male F2s. We found no evidence of main or interactive effects of maternal or paternal grandfather treatment for female F2 evasive swimming or freezing behaviors (Table 1; Figure 3). There was no correlation between the length of time frozen and whether or not individuals performed evasive swimming behaviors (t_247_=−0.58, p=0.56).

**Table 2:** Results of general linear mixed models testing predictors of F2 mass and length (n=125 males, n=126 females) and square-root transformed cortisol (n=97 males, n=99 females).

|  | Female F2s |  |  |  | Male F2s |  |  |  |
| --- | --- | --- | --- | --- | --- | --- | --- | --- |
|  | Mass |  |  |  |  |  |  |  |
|  | *t* | *df* | p | *Cohen's F* | *t* | *df* | p | *Cohen's F* |
| Maternal grandfather treatment | 0.83 | 18.84 | 0.42 | 0.08 | 0.82 | 14.26 | 0.43 | 0.08 |
| Paternal grandfather treatment | 1.88 | 16.36 | 0.08 | 0.05 | -1.35 | 117.21 | 0.18 | 0.13 |
| Standard length | 24.29 | 117.41 | **<0.001** | 2.34 | 24.31 | 114.33 | **<0.001** | 2.33 |
| Maternal * paternal GF treatment | -2.85 | 117.40 | **0.005** | 0.27 | --- | --- | --- | --- |
|  | Standard length |  |  |  |  |  |  |  |
|  | *t* | *df* | p | *Cohen's F* | *t* | *df* | p | *Cohen's F* |
| Maternal grandfather treatment | 1.22 | 5.58 | 0.27 | 0.12 | 0.94 | 117.10 | 0.35 | 0.09 |
| Paternal grandfather treatment | -0.002 | 2.69 | 1 | 0 | 0.56 | 14.96 | 0.59 | 0.05 |
| Tank density | -4.10 | 92.69 | **<0.001** | 0.41 | -4.86 | 107.67 | **<0.001** | 0.47 |
| Days since hatching | 0.27 | 70.63 | 0.27 | 0.03 | -1.33 | 59.33 | 0.19 | 0.13 |
|  | Acute cortisol |  |  |  |  |  |  |  |
|  | *t* | *df* | p | *Cohen's F* | *t* | *df* | p | *Cohen's F* |
| Maternal grandfather treatment | -0.58 | 7.87 | 0.58 | 0.06 | -0.04 | 10.62 | 0.97 | 0.004 |
| Paternal grandfather treatment | -0.43 | 7.15 | 0.68 | 0.05 | 0.12 | 10.75 | 0.90 | 0.02 |
| Standard length | -0.50 | 55.78 | 0.62 | 0.06 | 2.20 | 92.13 | **0.03** | 0.24 |

### Female F2s were heavier when their paternal grandfather, but not both grandfathers, was exposed to predation risk

For female F2s, we found an interaction between maternal grandfather and paternal grandfather treatment on mass (Table 2; Figure 4). Specifically, female F2s tended to be heavier when their paternal grandfather was exposed to predation risk (t_16.36_=1.88, p=0.08), but not when their maternal grandfather (t_18.84_=0.83, p=0.42) or both grandfathers (t_17.80_=−0.22, p=0.82) were exposed to predation risk. Model fits did not significantly worsen when maternal/maternal grandfather were removed (χ^2^=2.72, p=0.26), though they tended to worsen when random effects of paternal/paternal grandfather were removed (χ^2^=5.87, p=0.05). Male F2 mass was not altered by maternal or paternal grandfather treatment (Table 2; Figure 4). We found no evidence that stress-induced cortisol or length varied with paternal or maternal grandfather treatment for either male or female F2s (Table 2).

**Figure 4:**
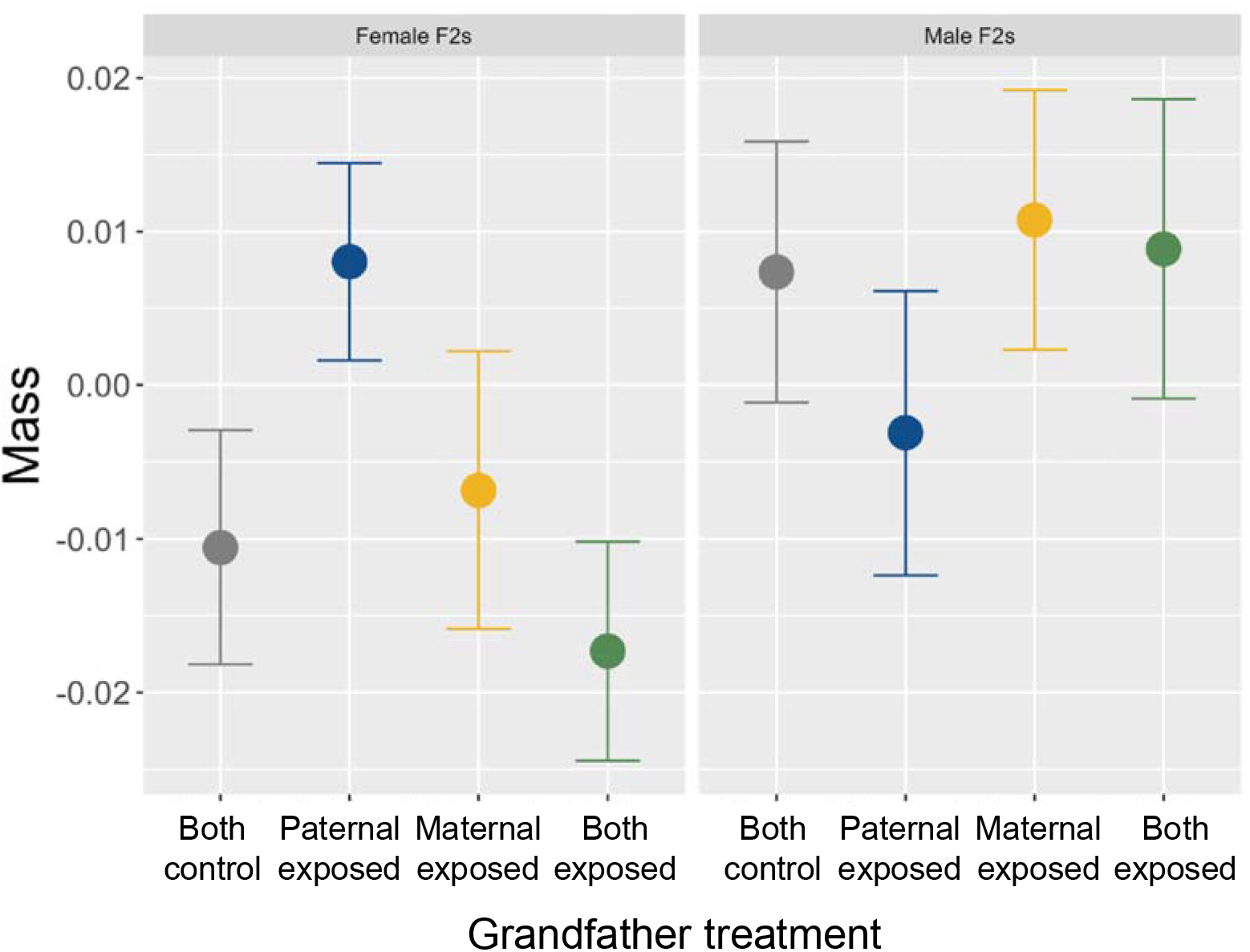
Mass (5 months post-hatching) of female (left; n=125) and male (right; n=123) F2s with two control grandfathers (grey), predator-exposed paternal grandfather (blue), predator-exposed maternal grandfather (yellow), or two predator-exposed grandfathers (green). Shown are the residuals of the regression model with grandpaternal treatment removed, plotted against grandpaternal treatment.

Although we found no effect of age on length, F2s were larger when they were in a lower density tank (Table 2). We found no detectable effect of size on activity/exploration or freezing behavior on either sex; while we found no detectable effect of length on stress-induced cortisol for females, larger male F2s had higher stress-induced cortisol (Table 1, 2).

## Discussion

In a previous study, we found that that paternal experience with predation risk had sexspecific consequences for male and female F1s (Hellmann *et al.* in review). These findings prompted the hypothesis that sex-specific effects in the F1 generation could contribute to sex and lineage-specific effects in the F2 generation. Here, we exposed F0 male sticklebacks to predation risk, reared the F1 generation in the absence of predation risk, and determined how the phenotypes of male and female F2s varied depending on whether predation risk was experienced by their paternal grandfather (inherited via F1 males), maternal grandfather (inherited via F1 females), or both grandfathers (inherited via both F1 males and females). We demonstrate that grandpaternal effects, mediated via sperm, are transmitted selectively to their grandoffspring. Specifically, female F2s were heavier and less responsive (reduced change in activity/exploration) to a simulated predator attack when their paternal grandfather was exposed to predation risk. In contrast, male F2s were less likely to perform evasive swimming (antipredator) behavior when their maternal grandfather was exposed to predation risk and tended to spend less time frozen when their maternal or paternal grandfather was exposed to predation risk. For all the above patterns, this change was present when one grandfather, but not both grandfathers, were exposed to predation risk. These findings suggest that grandpaternal effects are both sex-specific and lineage-specific: grandfathers’ experiences have different consequences for male and female F2s, and F2 traits depend on whether the paternal grandfather, maternal grandfather, or both grandfathers were exposed to predation risk. Consistent with recent metanalyses (Uller, Nakagawa & English 2013; Moore, Whiteman & Martin 2019; Yin *et al.* 2019), the effect sizes in this experiment were small to moderate.

We initially hypothesized that paternal transmission occurs along sex-specific lines (e.g. fathers to sons) but found little support for this hypothesis. Rather, we largely observed the opposite pattern, in which transmission was mediated across sexes from F1 males to F2 females and from F1 females to F2 males. Emborski and Mikheyev (2019) also found multigenerational transmission across sexes in fruit flies, whereby male offspring were influenced by the ancestral maternal diet and female offspring were influenced by the ancestral paternal diet. Further, a similar pattern of transmission to female descendants (F2s and F3s) via the paternal lineage has been documented in mammals in response to a wide range of maternal experiences, such as high-fat diets (Dunn & Bale 2011), chronic social instability (Saavedra-Rodríguez & Feig 2013), prenatal glucocorticoid exposure (Moisiadis *et al.* 2017) and food availability (Bygren *et al.* 2014), despite the fact that the cue originated in a different parent (F0 females versus males) and that the triggering cue varied across studies (e.g. diet versus predation risk).

These lineage effects may be generated by a number of different proximate mechanisms including genomic imprinting regulated in a sex-specific manner (Dunn & Bale 2011) or sexspecific embryonic responses to differences in sperm content (e.g. small RNAs). For example, an interesting possibility is that epigenetic changes to sex chromosomes are more stably transmitted via the F1 heterogametic sex (often males) due to lower rates of sex chromosome recombination (Bygren *et al.* 2014). From an ultimate perspective, sex-specific and lineage specific effects may evolve due to differences between males and females in 1) how the F1 generation perceive their environment and transmit cues of predation risk and 2) the different costs and benefits of attending to potentially outdated information from grandparents. Given this, future work examining how ecological divergence between males and females generates differences in predation exposure and perceived risk in the F0 and F1 generation in sticklebacks would be useful for understanding sex-specific transmission. Further, while our effects are seemingly independent of genetic variation within a population, evolved differences in antipredator genotypes across populations or species likely influence both exposure and perceived predation risk, thus potentially altering the strength and persistence of transgenerational effects (Walsh *et al.* 2016; Sentis *et al.* 2019) as well as the potential for sex-specific transmission.

Whether these phenotypic effects in the F2 generation are adaptive or non-adaptive is unclear. Previous studies have found that high predation environments can result in heavier offspring and earlier sexual maturity (Walsh & Reznick 2008; Mukherjee *et al.* 2014), suggesting that increased mass of female F2s could be adaptive. However, the reduced change in activity/exploration as well as lower levels of antipredator behavior suggest that F2s with predator-exposed grandfathers are actually less responsive to predation risk. This is consistent with the findings of Sentis *et al.* (2018) which found that, although aphid mothers exposed to predation risk produced many winged F1 morphs to facilitate dispersal, the frequency of winged morphs in the F2 generation was lower than the control group. It is currently unclear if this low antipredator defense in the F2 generation is adaptive (potentially driven by negative maternal effects (Kuijper & Hoyle 2015)) or non-adaptive. Indeed, it is quite possible that these sexspecific and lineage effects are not adaptive. Burke, Nakagawa and Bonduriansky (2019) suggest that complex sex-specific patterns may be particularly likely to be non-adaptive, because they reflect intralocus conflict that results from sex-specific selective pressures. These non-adaptive effects may be particularly likely to arise in species in which males are subject to strong sexual selection, as is the case in sticklebacks where there is high competition for breeding sites and females. Consequently, additional lineage-specific studies across a broader range of taxonomic groups, with diverse potential mechanisms of transmission, may determine the frequency of different patterns of transmission and whether these lineage-specific patterns of transmission are adaptive.

In addition to distinct grandpaternal effects via maternal and paternal lineages, we also found interactive effects: grandpaternal effects were evident if one grandfather was exposed to predation risk, but not if both grandfathers experienced predation risk. These interactive effects mean that if we had not isolated effects emerging in the paternal versus maternal lineage (e.g. compared controls to F2s with two predator-exposed grandfathers), we would have erroneously concluded that effects in the F1 generation did not persist until the F2 generation. This suggests that previous studies that have not examined these lineage effects may have underestimated the extent to which transgenerational environmental effects persist to the F2 generation. Interestingly, these lineage effects mirror the interactive effects between maternal and paternal cues that were observed in the F1 generation: offspring of predator-exposed fathers showed reduced survival against a sculpin predator, but this pattern was not evident when both the mother and father were exposed to predation risk (Hellmann *et al.* in review). This suggests the effects of paternal environments are moderated by maternal environments, both when mothers and fathers are directly exposed to the triggering environment (transmitted from predator-exposed F0s to the F1 generation) and when mothers and fathers inherit a cue about the environment from their parents (transmitted from the offspring of predator-exposed parents to the F2 generation). These interactive patterns may result because the combination of two parents/grandparents exposed to predation risk produces an entirely different phenotype compared to one parent/grandparent (Lehto & Tinghitella 2020; Hellmann *et al.* in review). Further, these non-additive interactions could resemble non-additive effects observed in genetic inheritance mechanisms, whereby phenomena such as the heterozygote advantage results in improved fitness of individuals with two different alleles relative to two matching alleles, particularly in fast-changing environments (Sellis *et al.* 2011).

An important finding emerging from this study, together with Hellmann *et al.* (in review), is that epigenetic transmission and phenotypic consequences can be decoupled: F2 antipredator behavior and mass were altered by grandpaternal exposure to predation risk, but not by paternal exposure to predation risk in the F1 generation. Similar results have been found in other systems (Panacek *et al.* 2011; Kim, Kwak & An 2013; Crocker & Hunter 2018), which collectively suggests that individuals may be silent carriers of epigenetic information, transmitting altered phenotypes to their offspring without actually displaying the phenotype themselves. Differences between phenotypes in the F1 and F2 generation may be linked to different proximate mechanisms that result in altered phenotypes in the F1 generations compared to F2 generations (Gapp *et al.* 2014; Bell & Hellmann 2019). For example, the F0 germline (sperm that will sire the F1 generation) is directly exposed to predation risk whereas the F1 germline is unexposed, potentially leading to different epigenetic changes in the F0 versus F1 sperm (Gapp *et al.* 2014). Alternatively, or in addition, cues from the F0 generation may alter how F1s experience their environment (e.g. social interactions (Giesing *et al.* 2011), habitat choice (Bestion *et al.* 2014)), which could induce additional epigenetic changes (Nagy & Turecki 2012) that are transmitted to the F2 generation and result in different phenotypes between the F1 and F2 generation. Future work examining differences and similarities in the mechanisms of transmission across multiple generations would be highly useful.

## Conclusions

Sex-specific and lineage-specific effects in externally-fertilizing organisms or organisms that lack parental care are not well characterized, as the majority of the studies on sex-specific and lineage-specific transgenerational plasticity have been carried out on maternal effects in mammals. Here, we demonstrate that similar sex-specific and lineage-specific effects can occur in different taxonomic groups (fish versus mammals), are observed when cues are transmitted via gametes alone (in the absence of *in utero* effects, parental care and differential allocation), and when cues are originally experienced by F0 males instead of F0 females. Together with Hellmann et al 2020, we demonstrate the need to consider biological sex when conducting transgenerational studies: male and female descendants often vary in their phenotypic response to parental experiences and complex non-additive patterns can arise from interactions between maternal and paternal cues. This selective inheritance has significant implications for theory, and raises questions regarding the mechanisms underlying the selective transmission of transgenerational information across generations and the interface between sex-specific selection pressures and the evolution of transgenerational plasticity. Sex-specific transgenerational plasticity could contribute to high degree of sexual dimorphism observed in many populations (Cooper, Gilman & Boughman 2011), especially if evolved differences in habitat use results in different levels of predation exposure for males and females, for instance. Future work examining interactions between female and male exposure (e.g., grandmaternal versus grandpaternal lineage effects) would help elucidate the ways in which sex-specific transgenerational mechanisms contribute to ecological divergence between males and females.

## Acknowledgements and funding

Thank you to Eunice Chen, Jack Deno, Erin Hsiao, Yangxue Ma, Raiza Singh, and Christian Zielinski for help with data collection and to the Bell lab for comments on previous versions of this manuscript. Thank you to Ryan Earley for help with the hormone assays. Thank you to Miles Bensky for creating the images for Figure 1 and Noelle James for the use of her photo in the graphical abstract. This work was supported by the National Institutes of Health award number 2R01GM082937-06A1 to Alison Bell and National Institutes of Health NRSA fellowship F32GM121033 to Jennifer Hellmann. There are no conflicts of interest for any of the authors.

## Authors contributions

JKH and AMB conceived of the study, JKH collected the data, ERC processed the cortisol samples, JKH analyzed the data and wrote the first draft of the manuscript, and JKH and AMB edited the manuscript.

## Data Accessibility

All datasets (behavior, morphology, cortisol) will be made available on Dryad upon acceptance of this manuscript.

