## Supplementary Material for "Sex-specific transgenerational plasticity II: Grandpaternal effects are lineage- and sex-specific in threespined sticklebacks"

***Clutch failure and density.*** Because it is possible that our experimental treatment resulted in differences in reproductive success in either the F0 or F1 generation, we evaluated clutch failure and clutch size for both generations. F0 paternal treatment did not alter fertilization rates or clutch survival in the F1 generation: 3/14 clutches of control parents and 2/13 clutches of predator-exposed fathers failed to survive to one month of age. F1 clutch size did not differ between control and predator-exposed fathers (n=22 clutches; mean control clutch size 27.9 ± 4.8 s.e., predator-exposed 28.0 ± 3.9 s.e.; unpaired t-test with unequal variance: t_19.11_=-0.01, p=0.99). Similarly, clutch failure in the F2 generation did not vary with treatment: 4/12 clutches with control grandfathers, 2/10 clutches with a predator-exposed paternal grandfather, 2/10 clutches with a predator-exposed maternal grandfather, and 2/10 clutches with two predator-exposed grandfathers failed to survive to one month of age. Clutch size was relatively consistent across F2 treatment groups (mean control: 20.3 ± 2.4 s.e., predator-exposed paternal grandfather: 16.3 ± 2.0 s.e., predator-exposed maternal grandfather: 19.6 ± 3.0 s.e., two predator-exposed grandfathers 18.5 ± 3.3 s.e.) and did not vary depending on the predation treatment of the maternal (t_27.58_=-0.31, p=0.76) or paternal (t_30.00_=0.98, p=0.34) grandfather.
